# Dynamics of bacterial biofilm development imaged using light sheet fluorescence microscopy

**DOI:** 10.1101/2024.07.10.602771

**Authors:** Lenka Šmerdová, Tibor Füzik, Lucie Valentová, Pavol Bárdy, Michaela Procházková, Martina Pařenicová, Pavel Plevka

**Affiliations:** Central European Institute of Technology, Masaryk University, Brno, Czech Republic; Veterinary Research Institute, Brno, Czech Republic

**Author notes:** Department of Chemistry, University of York, York, United Kingdom (Pavol Bárdy).

## Abstract

Biofilm formation exacerbates bacterial infections and interferes with industrial processes. However, the dynamics of biofilm development is not entirely understood. Here, we present a microfluidic cultivation system that enables continuous imaging of biofilm growth using light sheet fluorescence microscopy (LSFM). We studied the development of biofilms of the human pathogens *Staphylococcus aureus* and *Pseudomonas aeruginosa*. Due to the low phototoxicity of LSFM, biofilms can be continuously imaged without adverse effects on their development. Whereas *S. aureus* forms 50-70-μm-thick mushroom-like structures, a *P. aeruginosa* biofilm is 10-15 μm thick with cell clusters 25 μm in diameter. A combined biofilm, inoculated with an equal OD_600_ ratio of *S. aureus* and *P. aeruginosa*, resulted in the formation of large mushroom-like clusters of *S. aureus* cells that were subsequently dispersed by invading *P. aeruginosa.* A higher inoculation ratio favoring *P. aeruginosa* resulted in the formation of small and stable *S. aureus* clusters overgrown with *P. aeruginosa* cells. Applying conditioned media from *S. aureus* and *P. aeruginosa* coculture to a single-species *S. aureus* biofilm induced its dispersion. Integrating a microfluidic system into LSFM enables the visualization of biofilm formation dynamics and the effects of compounds on biofilm development.

## Introduction

*Staphylococcus aureus* is a human pathogen that causes skin, soft tissue, urinary tract, and implant-associated infections.^1, 2^ Biofilm formation is an essential aspect of *S. aureus* pathogenicity.^3^ Alongside *Pseudomonas aeruginosa, S. aureus* is a bacterial species implicated in respiratory tract infections, particularly in individuals with cystic fibrosis.^4^

Bacterial biofilms are organized aggregates of cells embedded in extracellular polymeric substances (EPS), including polysaccharides and DNA, and attached to a surface.^5^ Bacteria anchored in a biofilm are metabolically differentiated, and therefore some of them can resist antibiotics over extended periods.^6, 7^ Furthermore, the bulk size of the biofilm protects embedded bacteria from phagocytosis by immune cells.^8, 9^ Therefore, biofilm formation poses a therapeutic challenge.^10^

Biofilm formation is a complex process consisting of various stages that include attachment, proliferation, maturation, and dispersion.^11^ An important aspect of biofilm formation is the release of extracellular polymeric substances, a conglomeration of different biopolymers that form the scaffold of the three-dimensional biofilm structure. Members of the *Staphylococcus* genus produce robust mushroom-like biofilms.^12^ Extracellular polymeric substances released by *S. aureus* include polysaccharide intercellular adhesins (PIA), which mediate cell-to-cell adhesion and enable the attachment of cells to surfaces.^13–15^ In contrast, the initiation of biofilm formation by *P. aeruginosa* involves the release of Psl polysaccharides, but also the attachment of cells to surfaces by flagella and pili, indicating the critical role of bacterial motility in this process.^16, 17^ Extracellular DNA (eDNA) is another essential matrix component, originating from lysed cells.^18^ eDNA reinforces biofilm structure, interferes with host immune response, acts as a nutrient source, and serves as a gene pool for horizontal gene transfer.^19, 20^ Proteins with enzymatic activity in a biofilm matrix degrade exogenous macromolecules or EPS components to become available as nutrients during biofilm starvation. Structural proteins are involved in building the three-dimensional biofilm structure.^21^

Biofilm formation is a highly regulated process controlled by quorum-sensing systems that enable bacteria to sense population density.^22^ *S. aureus* employs an Agr quorum-sensing system, which regulates biofilm growth and dispersion.^23–25^ *P. aeruginosa* uses its quorum-sensing system during the early stages of biofilm development as a regulator of the production of virulence factors.^26^

*P. aeruginosa* is frequently recognized as a co-colonizer along with other bacterial species. Co-infection of the lungs of cystic fibrosis patients by *P. aeruginosa* and *S. aureus* is an indication of poor clinical outcome.^4^ Bacteria in mixed-species biofilms can compete for resources and space or benefit from each other.^27^ The relationship between *P. aeruginosa* and *S. aureus* is complex, and the impact of their coexistence in polymicrobial infections remains challenging to predict.

Here, we present an imaging system that enables continuous observation of biofilm development. The system combines a light sheet fluorescent microscope (LSFM) with a microfluidic system to monitor bacterial growth. The main advantages over the other available approaches are three-dimensional imaging capabilities, high spatial and temporal resolution, and low phototoxicity.^28^ LSFM is unique in providing a relatively large imaging area.^28–30^ Our system enables the administration of selected substances to biofilms and subsequent observation of their impact on its development.

## Results and discussion

### Continuous imaging of biofilm development by LSFM

Biofilm development is a complex process spanning multiple days during which the biofilm-forming cells need to be supplied with nutrients.^31^ Imaging biofilm development, using most of the currently available fluorescent microscopy techniques, is challenging because the biofilm-forming cells become exposed to intense light, and the resulting phototoxicity disrupts the natural processes. The predominantly used confocal imaging enables a few snapshots of biofilm development to be recorded.^32, 33^ To permit continuous biofilm imaging, we constructed an imaging setup consisting of an LSFM and a microfluidic cell supplied with fresh medium by a smooth flow pump (Fig. 1). LSFM is a planar illumination technique in which the axis of illumination is perpendicular to the optical axis of the detection objective.^28^ The imaging system of the Zeiss Z.1 Lightsheet microscope used in our setup combines a two-sided light sheet illumination with an orthogonal detection objective. The laser light shaped into a “sheet” only excites fluorophores within the focal volume of the detection objective, resulting in a reduction in photobleaching and phototoxicity compared to other fluorescence imaging approaches.^28–30^ Therefore, LSFM is particularly suited for the long-term imaging of living cells.

**Fig. 1.**
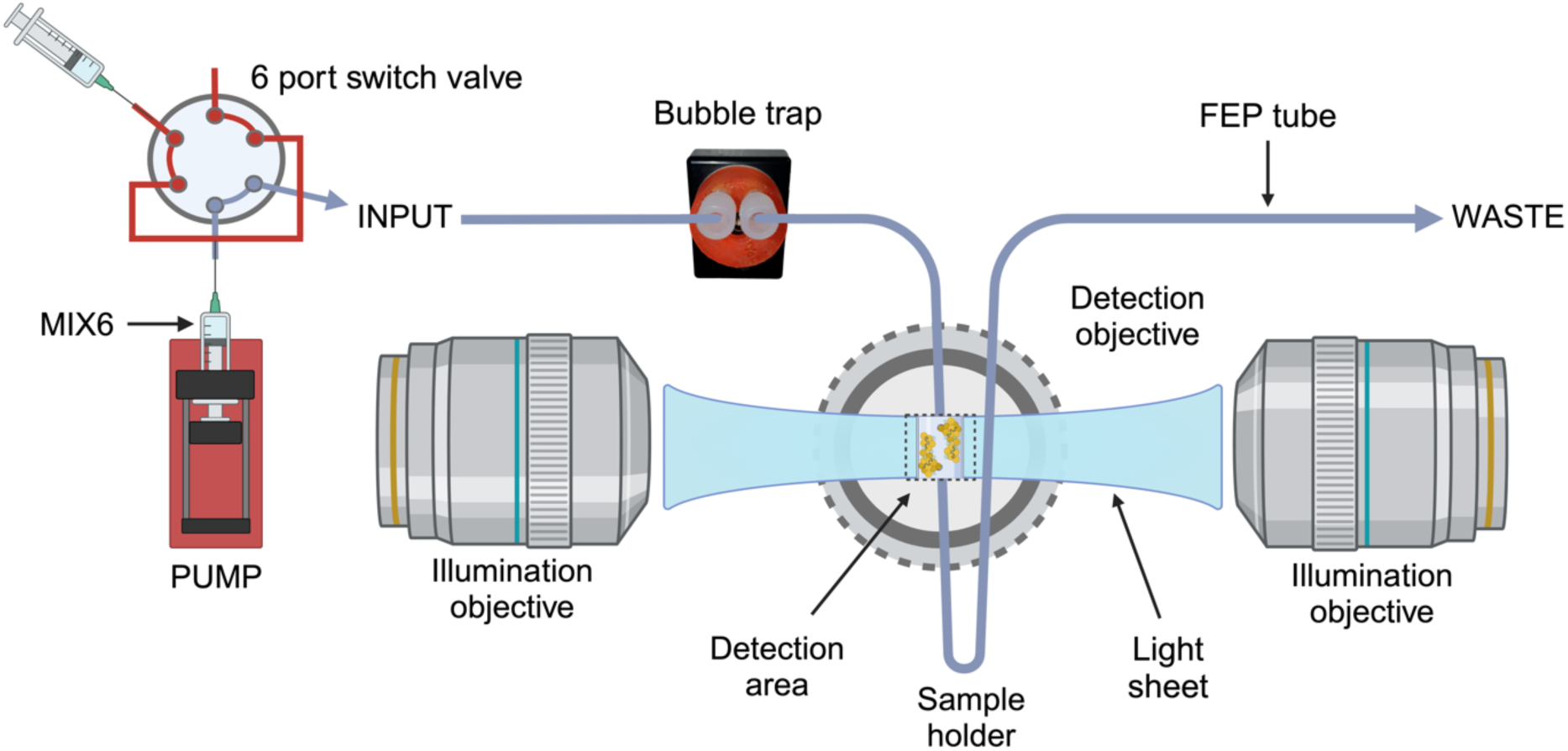
Scheme of LSFM imaging of a biofilm growing on the inner wall of an FEP tube. A syringe pump drives the flow of fresh medium to the biofilm.

In our system, biofilms are cultivated at the inner face of a fluorinated ethylene-propylene (FEP) tube with an inner diameter of 300 μm (Fig. 1). The tube is positioned in the imaging chamber of the LSFM microscope, which is filled with water kept at 37°C. FEP is optically clear and has a refractive index of 1.344, similar to that of water (1.335), resulting in a minimal bending of light at tube-water interfaces. Furthermore, the tubing made of FEP is sturdy but flexible. To enable the efficient detection of fluorescent signals, we developed a clear medium with negligible background fluorescence called MIX6. The medium flow through the biofilm-containing tubing is driven by a smooth flow pump (Fig. 1). We use a pumping rate that produces a speed of liquid movement through the FEP tube of 11 μm/s, resulting in a flow rate of 2.7 μl/hour. The flow at this speed is laminar and sufficient to supply fresh medium to the attached cells while not damaging the biofilm structure. The microfluidic system includes a two-position six-port switching valve, which enables the administration of additional substances through the medium flowing to the observed section of a biofilm. A bubble trap ensures that accidental air bubbles introduced into the system do not destroy the biofilm structure.

The motorized sample holder stage of the Zeiss Z.1 Lightsheet microscope enables the multidirectional imaging of samples by rotating them in the imaging chamber. Images of a sample recorded from multiple directions can be combined to produce a three-dimensional reconstruction.^34^ Due to the tubing bringing fresh medium to the imaged cells, the rotation of the sample in our setup is limited to 180°, which enables the recording of information necessary to calculate a reconstruction of the imaged volume with isotropic resolution.^35^ In our setup, the thickness of the excitation light sheet is 3.8 μm at the focal point. The depth of focus of the imaging objective is 1,52 μm, and we used a 0.53 μm spacing of frames in the direction perpendicular to the imaging plane. The numerical aperture of the imaging objective is 1.0, and we set the imaging area to 373 μm x 373 μm sampled to 1920 x 1920 pixels, resulting in a pixel size corresponding to 0.194 μm.

The three-dimensional reconstructions of the imaged volumes were calculated using the BigStitcher plugin in the software package FIJI.^34, 35^ The reconstruction process consists of interest point detection and registration, alignment of image stacks, image fusion, and deconvolution.^35^ We utilized the signal from bacteria expressing fluorescent proteins to serve as interest points for the alignment.

### Structure and dynamics of *S. aureus* biofilm development

The formation of *S. aureus* biofilms is a multi-step process that has been shown to include cell attachment, multiplication, exodus, maturation, and dispersal^8, 12^. To visualize the biofilm development, we labeled *S. aureus* cells by expression of the red fluorescent protein mCherry2-L under the control of the *S. aureus* constitutive *capA* promoter.^36^ In the *S. aureus* genome, the promoter controls the synthesis of capsular polysaccharides, which play a role in adherence and pathogenesis.^37, 38^ The expression of mCherry2-L under the control of the capA promoter positively correlates with the metabolic activity of the labeled cells.^37^ The fluorescent dyes POPO-1 and WGA Alexa Fluor 488 were included in MIX6 media to label extracellular DNA and PIA (poly-β(1-6)-N-acetylglucosamine), respectively. Planktonic *S. aureus* cells expressing mCherry2-L were injected into an FEP tube and allowed to adhere to its inner wall for 4 hours. The floating cells were removed by pumping fresh medium through the tube. The biofilm growth was imaged every 30 min for 48 hours (Fig. 2, Supplementary movie 1).

**Fig. 2.**
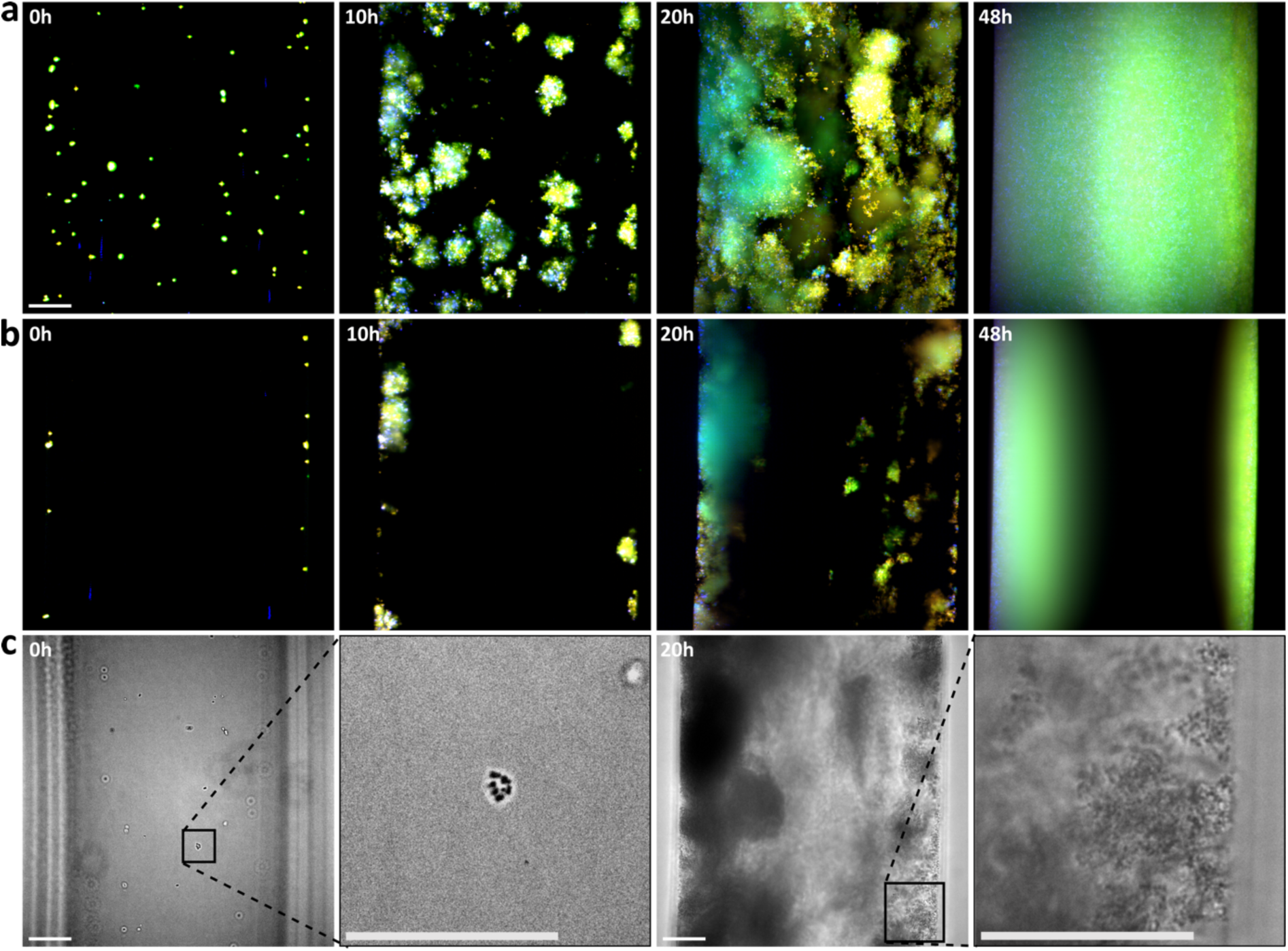
Time-lapse imaging of *S. aureus* biofilm development. *S. aureus* cells are shown in orange, eDNA in blue, and PIA in green. **(a)** Maximum intensity projections of *S. aureus* biofilms growing on the inner face of an FEP tube with an inner diameter of 300 μm. **(b)** Maximum intensity projection of 53-μm-thick central part of the FEP tube. **(c)** Images of the FEP tube with attached *S. aureus* cells recorded in visible light. Scale bars 50 μm.

Immediately after attachment, *S. aureus* cells started to produce an extracellular matrix (Fig. 2 a,b), which confirms the importance of PIA and eDNA for the initial steps of colonization of a surface and microcolony formation.^5, 39^ After 10 hours of development, *S. aureus* cells formed clusters that were 30-40 μm in size (Fig. 2 a,b). After 20-30 hours, *S. aureus* formed a 50-70-μm-thick layer that covered most of the available surface (Fig. 2, 3). Individual *S. aureus* cells continuously detached from the biofilm matrix during biofilm growth. The cell release was preceded by an increasing release of eDNA (Supplementary Movie 1), indicating increased cell death. Cell lysis results in the release of enzymes that degrade the biofilm matrix.^40, 41^

The three-dimensional multiview reconstruction of a 20-hour-old *S. aureus* biofilm was of sufficient quality to distinguish individual cells, clusters of extracellular DNA, and the diffuse distribution of PIA (Fig. 3). At this stage of biofilm development, *S. aureus* cells are distributed throughout the biofilm volume. Cells close to the FEP tube exhibit higher fluorescence than those in the other parts of the biofilm (Fig. 3 g,k, Supplementary Fig. 1). The more intense signal of *S. aureus* cells corresponds to increased metabolic activity. The high degree of cell heterogeneity within the biofilm is a well-known phenomenon of *S. aureus* biofilms, including fast-growing, slow-growing, and cells in stationary or dormant phases.^7^ The distribution of metabolically active cells changed as the biofilm matured, with more metabolically active cells located at the surface of the biofilm exposed to the medium (Supplementary Fig. 2). Cells located in these regions of the biofilm continued to be metabolically active, probably due to the higher availability of nutrients and oxygen than in the deeper parts of the biofilm.^42^ The distribution of eDNA followed the distribution of *S. aureus* cells, consistent with its origin in cell lysis and the role of eDNA in biofilm architecture.^19^ PIAs were predominantly in the middle of biofilm clusters (Fig. 3 e-l, Supplementary Fig. 1), anchoring staphylococcal cells to the biofilm matrix.^43^

**Fig. 3.**
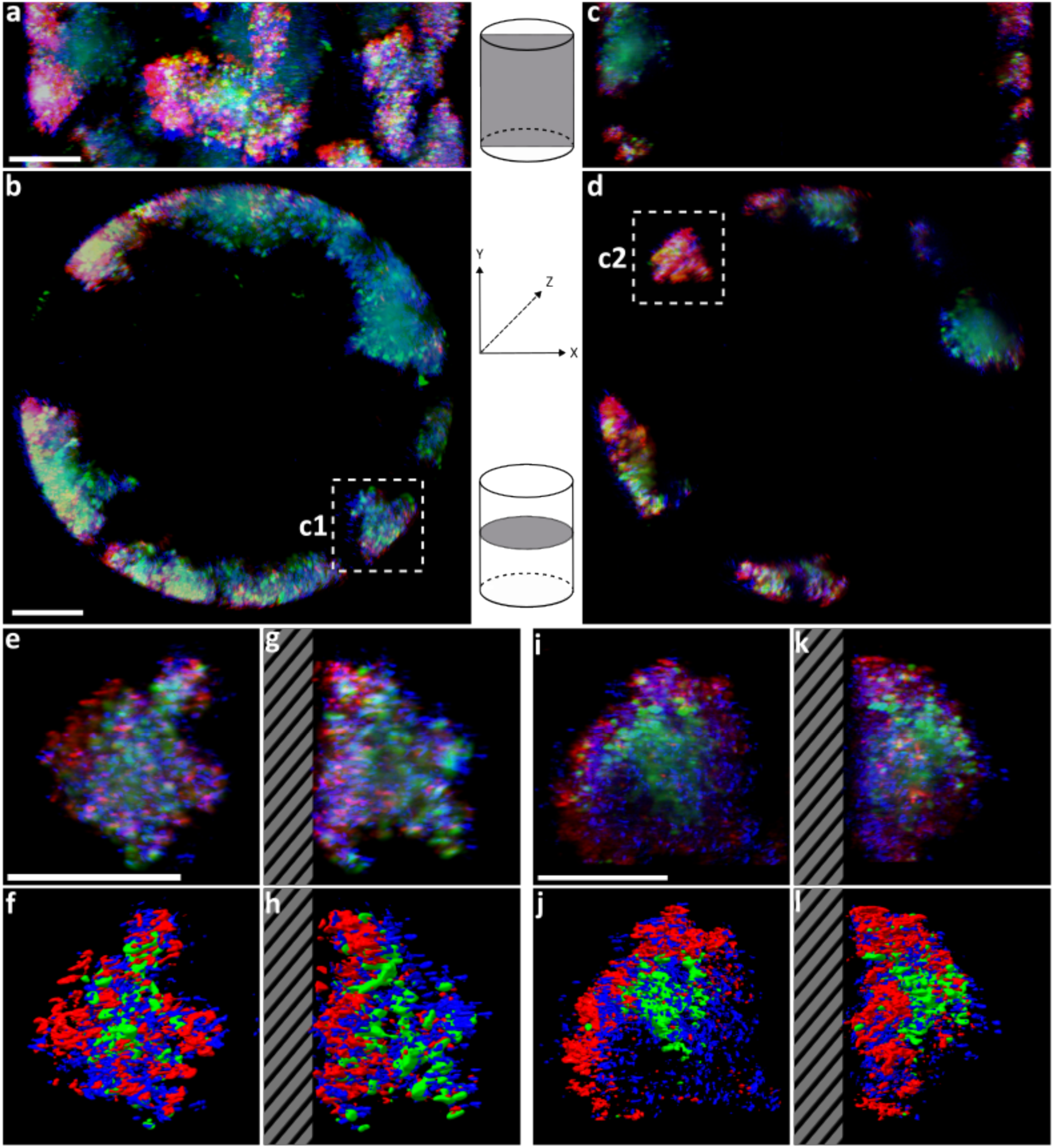
Three-dimensional reconstruction of 20-hour-old *S. aureus* biofilm. *S. aureus* cells are shown in orange, eDNA in blue, and PIA in green. **(a,b)** Projections of a three-dimensional reconstruction of an FEP tube segment containing an *S. aureus* biofilm. The direction of the projection is along the FEP tube axis (a) and perpendicular to the FEP tube axis (b). **(c,d)** Projection of a 6-μm-thick central section through the reconstruction of the biofilm-containing FEP tube in a direction along (c) and perpendicular (d) to its axis. The insets in panels b and d indicate the positions of biofilm clusters, which are shown in detail in panels e-l. Images of clusters c1 **(e-h)** and c2 **(i-l)**. Projections of the biofilm clusters in a direction tangential (e, i) and perpendicular (g, k) to the FEP tube wall. Surface representations of the biofilm reconstructions are displayed using ChimeraX **(f,h,j,l)**. Dashed areas in panels (g,h,k,l) indicate cross sections of the wall of an FEP tube. Scale bars 50 μm.

### Structure and dynamics of *P. aeruginosa* biofilm development

To visualize *P. aeruginosa* biofilm development, the cells were transformed with a plasmid expressing fast-maturating superfolder GFP (msfGFP) ^44^ under the *XylS/Pm* promoter induced by the addition of 1 mM 3-methyl benzoate (3-Mbz).^45^ Extracellular DNA was labelled using BOBO-3 stain. *P. aeruginosa* cells were allowed to attach to the inner surface of an FEP tube using the same approach as that described above for *S. aureus,* and the biofilm growth was imaged every 30 minutes for 48 hours (Fig. 4, Supplementary Movie 2). Some *P. aeruginosa* cells lysed immediately after attachment and released their DNA (Fig. 4 a, b, Supplementary Movie 2). The release of the DNA can promote the adhesion of other cells and lead to cell aggregation.^46^ It was shown previously that a subpopulation of cells in a *P. aeruginosa* biofilm undergo cell lysis accompanied by the liberation of cytosolic content that can serve as nutrients for other cells and boost their growth.^47^ Thirteen hours after attachment, *P. aeruginosa* cells proliferated extensively (Supplementary movie 2). After 20 hours, *P. aeruginosa* formed a 10-15-μm-thick biofilm covering the FEP tube’s inner wall with cell clusters ranging from 20-25 μm in diameter (Fig. 4, 5). It has been shown that the structure of a *P. aeruginosa* biofilm is affected by the surface motility of the cells.^48^ Flat, uniform biofilms are thought to be associated with a high surface motility of cells, whereas biofilms containing cell aggregates are related to the limited movement and clonal growth of attached cells.^48^

**Fig. 4.**
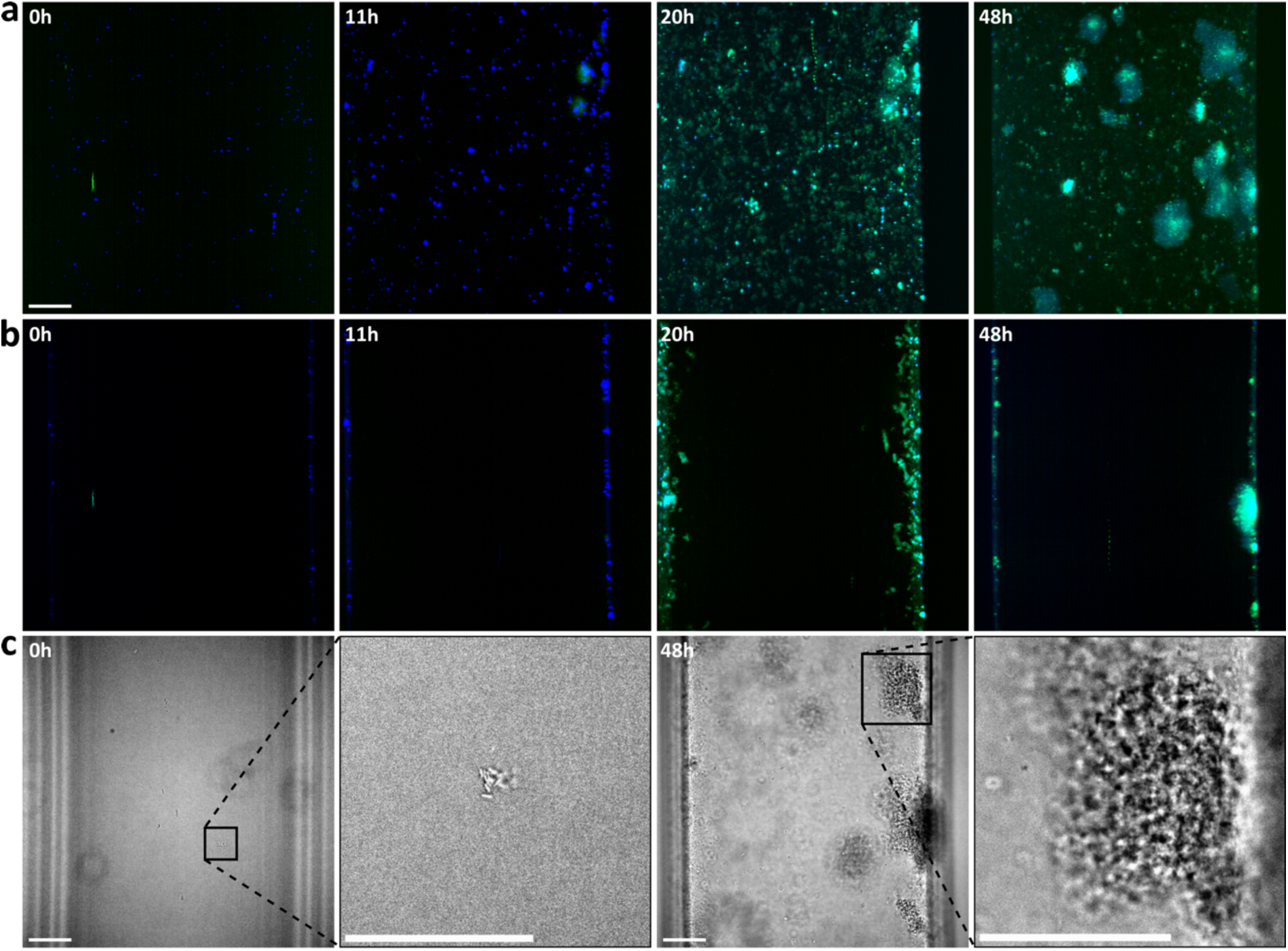
Time-lapse imaging of *P. aeruginosa* biofilm development. *P. aeruginosa* cells labeled by expression of msfGFP are shown in green and eDNA in blue. **(a)** Maximum intensity projections of *P. aeruginosa* biofilms growing at the inner face of an FEP tube with an inner diameter of 300 μm. **(b)** Maximum intensity projection of 53-μm-thick central part of the FEP tube. **(c)** Images of the FEP tube with attached *P. aeruginosa* cells recorded in visible light. Scale bars 50 μm.

### Development and organization of combined *P. aeruginosa* and *S. aureus* biofilm depend on relative species abundance

*P. aeruginosa* and *S. aureus* are the most prevalent pathogens identified in the wounds and sputum of cystic fibrosis patients.^49^ ^50^ Co-infection by *P. aeruginosa* and *S. aureus* leads to a reduced sensitivity of both bacterial species to antibiotics.^51^ It is therefore essential to understand the development of the combined biofilms. Here, we studied the development of combined biofilms initiated by mixed planktonic cultures with an equal OD_600_ ratio of *P. aeruginosa* and *S. aureus* cells and a mixture with a five times higher relative concentration of *P. aeruginosa*.

To initiate the formation of a mixed-species biofilm, FEP tubing was inoculated with equal amounts of fluorescently tagged *P. aeruginosa* and *S. aureus* (OD_600_ 0.01, which correspond to 6.7×10^6^ CFU/ml for *S. aureus* and 4.1 x 10^7^ CFU/ml for *P. aeruginosa*) (Fig. 6a, Supplementary movie 3). *S. aureus* proliferated faster than *P. aeruginosa* in the initial stages of mixed-species biofilm development (Fig. 6, Supplementary Movie 3, Supplementary Fig. 3). The abundance of *P. aeruginosa* increased after 9-12 hours of biofilm growth (Fig. 6 a). *S. aureus* formed 40-50-μm-sized clusters, which became gradually overgrown on the surface with *P. aeruginosa* cells (Fig. 6a). After 22 hours of development, *S. aureus* biofilm clusters gradually disintegrated (Fig. 6a, Supplementary movie 3). The final stage of the combined biofilm seeded by equal OD_600_ ratios of *P. aeruginosa* and *S. aureus* was a thin layer covering the FEP surface, which contained predominantly *P. aeruginosa* cells (Fig. 6 a, b). The development of a *P. aeruginosa* and *S. aureus* mixed biofilm was different when the culture was initiated with a cell mixture containing a higher amount of *P. aeruginosa* (OD_600_ 0.05) than that of *S. aureus* (OD_600_ 0.01). Under these conditions, more *P. aeruginosa* cells colonized the FEP surface and started to proliferate three hours after cell attachment (Supplementary movie 4). The size of *S. aureus* colonies was limited to 20-25 μm, probably due to the presence of abundant *P. aeruginosa* cells (Fig. 6 c). Despite the predominance of *P. aeruginosa,* which completely covered the surface, the smaller *S. aureus* colonies remained stable and did not disintegrate (Fig. 6 c, 7, Supplementary movie 4). This phenotype resembles the well-documented small colony variant (SCV) phenotype of *S. aureus,* which is related to a metabolic switch from oxidative to fermentative metabolism induced by respiratory toxins released by *P. aeruginosa.*^52^ After 64 hours of *P. aeruginosa* and *S. aureus* biofilm growth, initiated by 5 : 1 ratio of cultures in favor of *P. aeruginosa*, *S. aureus* clusters no longer grew and remained covered with a thin layer of *P. aeruginosa* (Fig. 7). It has been shown that *P. aeruginosa* can outgrow *S. aureus* under some conditions, but the bacteria can coexist and benefit from each other.^27^ Interactions of *P. aeruginosa* and *S. aureus* in a biofilm range from initial competition through adaptation to cooperation.^53^

**Fig. 5.**
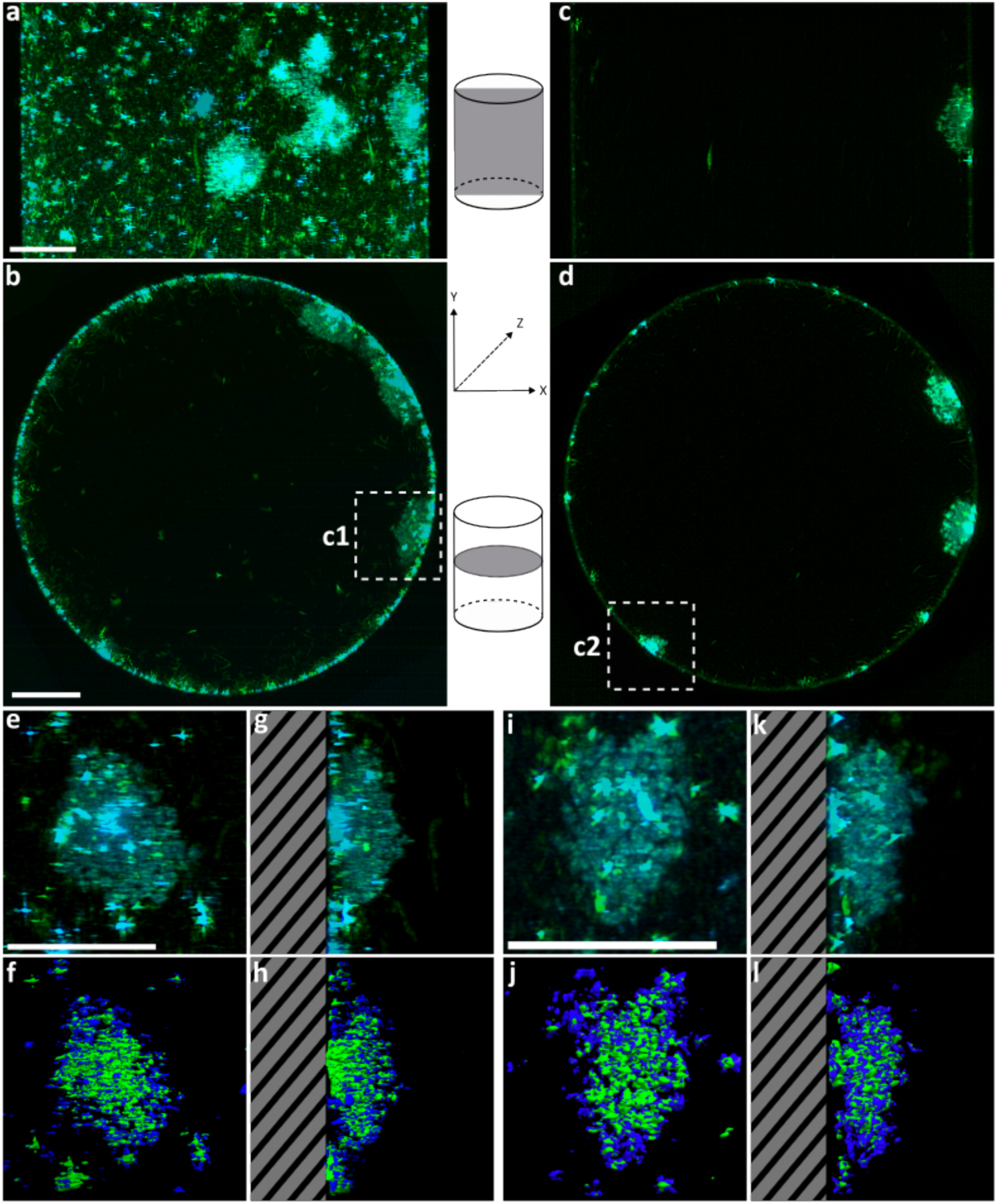
Three-dimensional reconstruction of 23-hour-old *P. aeruginosa* biofilm. *P. aeruginosa* cells are shown in green, and eDNA in blue. **(a,b)** Projections of a three-dimensional reconstruction of an FEP tube segment containing *P. aeruginosa* biofilm. The direction of the projection is along the FEP tube axis (a) and perpendicular to the FEP tube axis (b). **(c,d)** Projection of a 6-μm-thick central section through the reconstruction of the biofilm-containing FEP tube in a direction along (c) and perpendicular (d) to its axis. The insets in panels b and d indicate the positions of biofilm clusters, which are shown in detail in panels e-l. Images of clusters c1 **(e-h)** and c2 in panel d **(i-l)**. Projections of the biofilm clusters in a direction tangential (e, i) and perpendicular (g, k) to the FEP tube wall. Surface representations of the biofilm reconstructions are displayed using ChimeraX **(f,h,j,l)**. Dashed areas in panels (g,h,k,l) indicate cross sections of the wall of an FEP tube. Scale bars 50 μm.

**Fig. 6.**
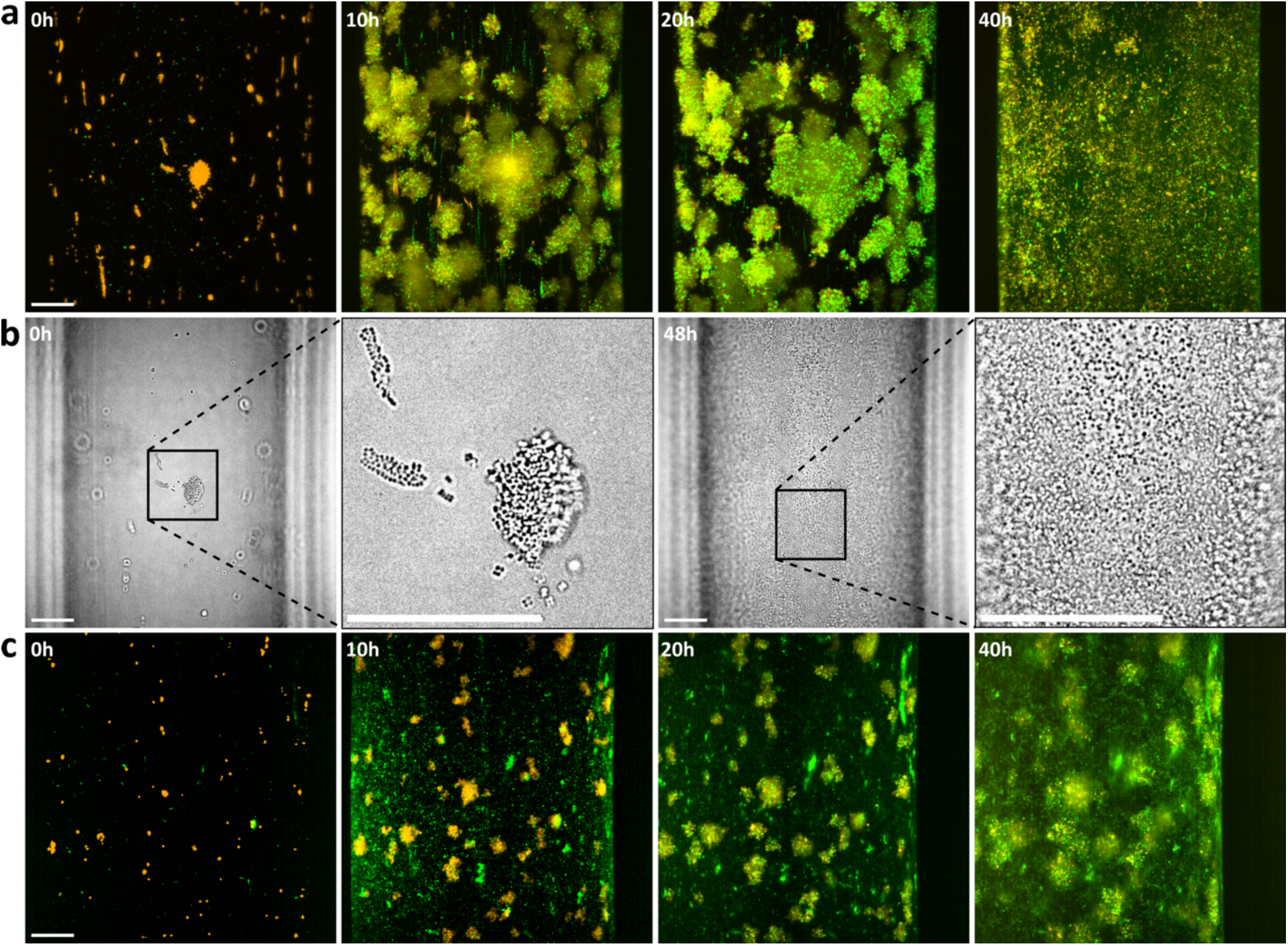
Time-lapse of the development of mixed *P. aeruginosa* and *S. aureus* biofilms. *P. aeruginosa* cells are shown in green, and those of *S. aureus* in orange. **(a)** Coculture inoculated using equal OD_600_ inoculation ratio. Maximum intensity projections of mixed biofilms growing at the inner face of an FEP tube with an inner diameter of 300 μm. **(b)** Coculture inoculated using equal OD_600_ inoculation ratio. Images of the FEP tube with attached cells recorded in visible light. **(c)** Higher OD_600_ inoculation ratio in favor of *P. aeruginosa* (5:1). Scale bars 50 μm.

**Fig. 7.**
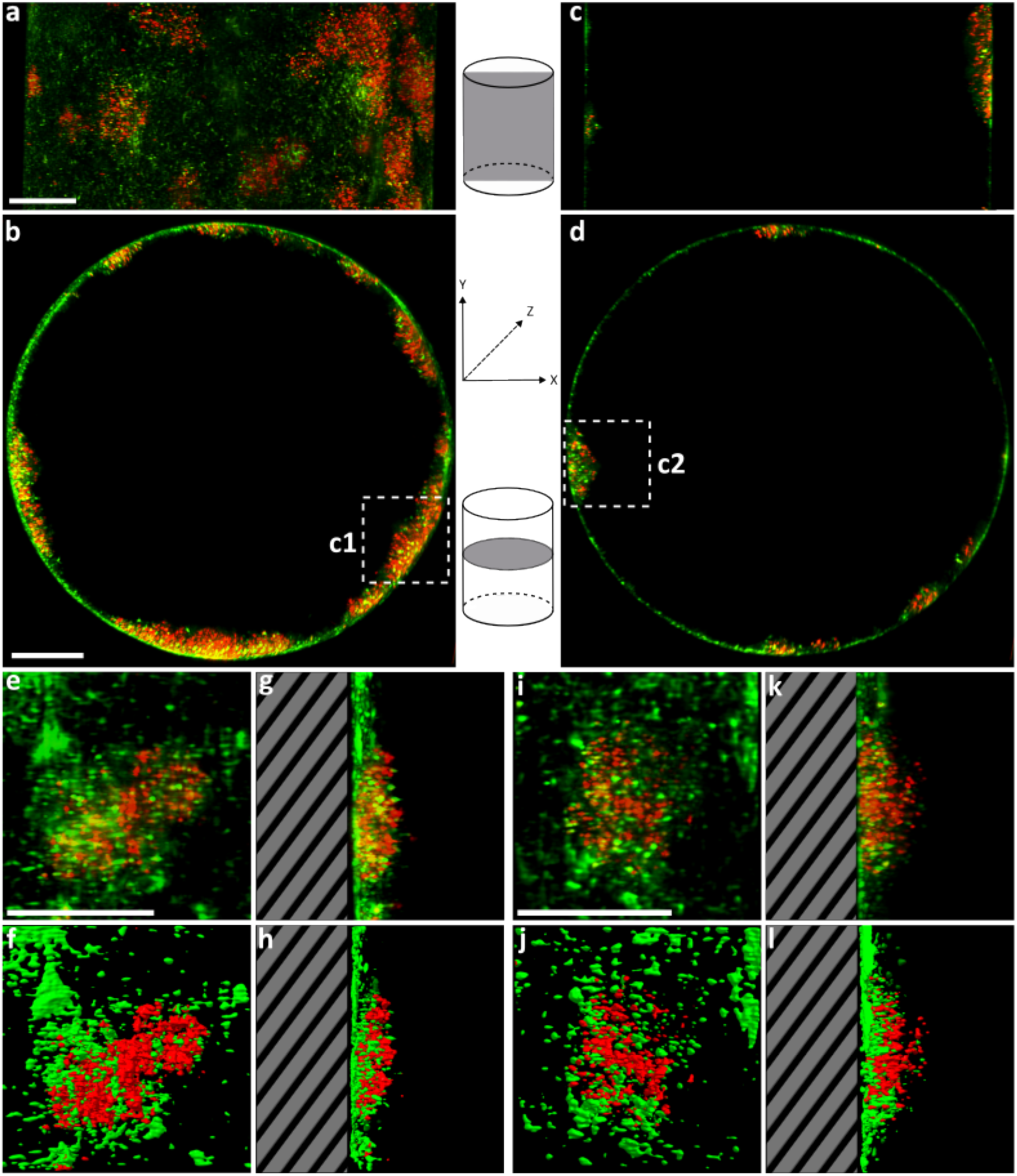
Three-dimensional reconstruction of mixed *P. aeruginosa* and *S. aureus* biofilm after 64 hours of development. *P. aeruginosa* cells are shown in green, and *S. aureus* cells are shown in red. The coculture was inoculated using a 5:1 OD_600_ ratio in favor of *P. aeruginosa*. **(a,b)** Projections of a three-dimensional reconstruction of an FEP tube segment containing a mixed biofilm. The direction of the projection is along the FEP tube axis (a) and perpendicular to the FEP tube axis (b). **(c,d)** Projection of a 6-μm-thick central section through the reconstruction of the biofilm-containing FEP tube in a direction along (c) and perpendicular (d) to its axis. The insets in panels b and d indicate the positions of biofilm clusters, which are shown in detail in panels e-l. Images of clusters c1 **(e-h)** and c2 in panel d **(i-l)**. Projections of the biofilm clusters in a direction tangential (e, i) and perpendicular (g, k) to the FEP tube wall. Surface representations of the biofilm reconstructions are displayed using ChimeraX **(f,h,j,l)**. Dashed areas in panels (g,h,k,l) indicate cross sections of the wall of an FEP tube. Scale bars 50 μm.

### Metabolites produced by *P. aeruginosa* induce dispersion of *S. aureus* biofilm

*P. aeruginosa* produces compounds, including respiratory toxins (quinoline N-oxides, pyocyanin, hydrogen cyanide)^54^, LasA proteases that lyse *S. aureus* cells, and rhamnolipids and Cis-2-decenoic acid, which can trigger *S. aureus* biofilm dispersal.^55^ The production of these compounds is regulated by quorum sensing (QS) and *Pseudomonas* quinolone signaling (PQS) systems.^27, 56, 57^ To study the effect of metabolites produced by *P. aeruginosa* on a *S. aureus* biofilm, we prepared a conditioned medium from the static co-cultivation of *P. aeruginosa* and *S. aureus*. The conditioned medium has blue background fluorescence, indicating the presence of secondary metabolites (Fig. 8). *S. aureus* cells cultivated in the conditioned medium did not start to proliferate even after 20 hours (Fig. 8a). After 21 hours of cultivation, the conditioned medium was replaced with fresh MIX6 medium; however, *S. aureus* cells did not start to divide within the 20-hour observation period, which indicates a long-term inhibition of proliferation caused by *P. aeruginosa* metabolites (Fig. 8 a).

**Fig. 8.**
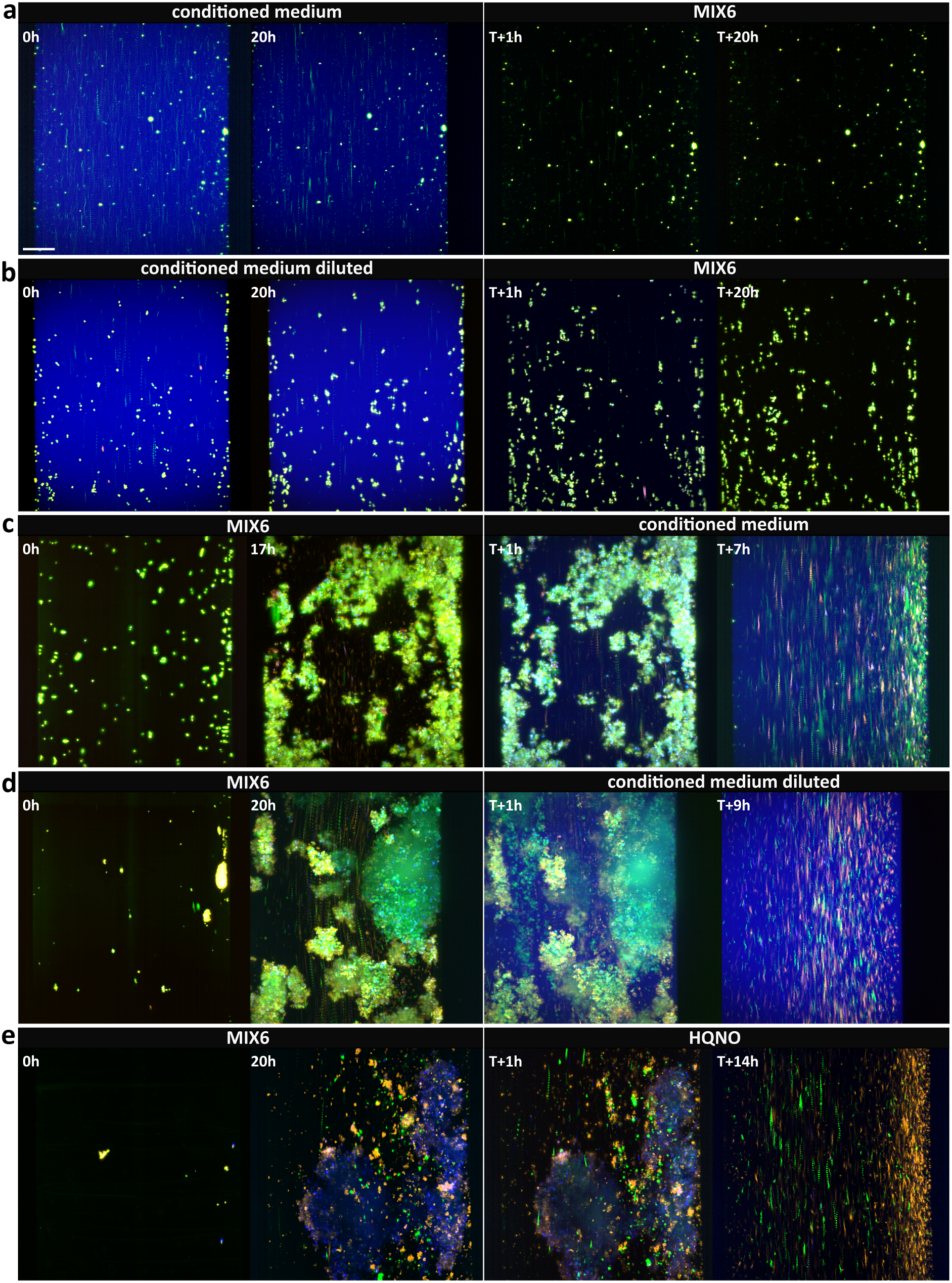
*S. aureus* biofilm development in conditioned medium. *S. aureus* cells are shown in orange, eDNA in blue, and PIA in green. All images represent maximum intensities projections composed from all slices of the FEP tube. **(a)** *S. aureus* cultivated in a conditioned medium. Timepoint T marks the replacement of the conditioned medium with MIX6 medium. **(b)** *S. aureus* was cultivated in a conditioned medium diluted 1:1 with MIX6 medium. Timepoint T marks the replacement time of the diluted conditioned medium with MIX6 medium. **(c)** *S. aureus* cultivated in MIX6 medium. Timepoint T marks the replacement of MIX6 medium with conditioned medium. **(d)** *S. aureus* cultivated in MIX6 medium. Timepoint T marks the replacement of MIX6 medium with diluted conditioned medium. **(e)** *S. aureus* cultivated in MIX6 medium. Timepoint T marks the addition of HQNO 150 ng/ml to the MIX6 medium. Scale bars 50 μm.

When added to a pre-formed 18-hour-old *S. aureus* biofilm, the conditioned medium induced its dispersal (Fig. 8 c). The biofilm-embedded *S. aureus* cells stopped dividing after immersion into the conditioned medium and dispersed after 7 hours (Fig. 8 c, Supplementary movie 5). To differentiate the effect of compounds released by *P. aeruginosa* from the effects of nutrient depletion, which may be due to the previous cultivation of a mixed biofilm in the conditioned medium, the experiments were repeated with a conditioned medium diluted with fresh MIX6 in a 1:1 ratio (Fig. 8 b, d, Supplementary movie 5). The effects on *S. aureus* biofilm formation and the induction of its dispersal were the same as those of the undiluted conditioned medium (Fig. 8 b, d).

*P. aeruginosa* produces an array of secondary metabolites that interfere with the growth of other bacterial species, including *S. aureus*. LC-MS/MS analysis identified the presence of respiratory toxins in conditioned media produced by *P. aeruginosa S. aureus* coculture (Supplementary Fig. 4). The media contained exoproducts of the *Pseudomonas* quinolone signaling pathway NQNO and C9, which target the respiratory electron transport chain and suppress the growth of *S. aureus*.^54^ The concentration of respiratory toxins released by *P. aeruginosa* in a single-species culture was 1.5-2 times higher than that from *S. aureus* and *P. aeruginosa* coculture (Supplementary Fig. 4). These results indicate that *P. aeruginosa* reacts to *S. aureus* cells in a coculture, contributing to the mutually regulated coexistence of these pathogens, as was shown previously for the coculture of *P. aeruginosa* and *Klebsiella pneumoniae*.^58^ Here, we show that the *S. aureus* biofilm disintegrated after 14 hours of cultivation in media containing 150 ng/ml HQNO, the concentration produced by *P. aeruginosa* in the coculture. This is consistent with quinoline N-oxides suppressing the growth of *S. aureus* (Fig. 8 e, Supplementary movie 6).

The conditioned media from *P. aeruginosa S. aureus* coculture contained surfactants (Supplementary Fig. 5)^59^. It has been shown that *P. aeruginosa* secretes rhamnolipids that promote *S. aureus* biofilm dispersal.^56^ These QS-regulated biosurfactants reduce surface tension and can permeabilize the membranes of microorganisms, resulting in the leakage of intracellular material.^55^

## Conclusions

We developed a system combining LSFM with a microfluidic cultivation of bacteria inside an FEP tube to enable the continuous imaging of biofilm development. The system can be used to visualize the interactions between bacteria in a coculture and monitor the effects of various substances on biofilm development. Multidirectional LSFM data can be used to calculate three-dimensional reconstructions of biofilm structure by multiview registration.^35^ The three-dimensional reconstructions make it possible to determine the distribution of cells and extracellular matrix components in a biofilm. We show that the type of biofilm formed by *S. aureus* and *P. aeruginosa* coculture is determined by the initial ratio of cells that seed a surface. *S. aureus* and *P. aeruginosa* coexistence can result in a degradation of *S. aureus* biofilm or the coexistence of the two bacterial species in a combined consortium. The microfluidic system can be used to study the effects of the conditioned medium or specific substances on biofilm development. Respiratory toxins secreted by *P. aeruginosa* cause *S. aureus* biofilm dispersion. LSFM, with an integrated microfluidic system, represents a powerful tool for studying biofilm formation and investigating the effects of antimicrobial agents.

## Materials and Methods

### Bacterial strains and media

The bacterial strains and plasmids used in this study are listed in Table S1. Overnight cultures of *S. aureus* were cultivated at 37°C in meat peptone medium: nutrient broth 13 g/L (Thermo Fisher Scientific); yeast extract 3 g/L (Carl Roth); peptone 5 g/L (Thermo Fisher Scientific). *P. aeruginosa* was cultivated in Luria-Bertani medium (Sigma-Aldrich). MIX6 medium was used to cultivate bacteria for LSFM imaging. MIX6 medium consists of 40 mM Na_2_HPO_4_ (Sigma-Aldrich); 22 mM KH_2_PO_4_ (Carl Roth); 9.35 mM NH_4_Cl (Penta Chemicals); 0.2 mM MgSO_4_ (Sigma-Aldrich); 0.2% glucose (Penta Chemicals); 5 ml/L MEM vitamin solution (Thermo Fisher Scientific); 150 ml/L MEM non-essential amino acid solution (Sigma-Aldrich); 100 ml/L MEM essential amino acid solution (Sigma-Aldrich); 0.1 mg/L biotin (Sigma-Aldrich); 0.85 g/L L-proline (Sigma-Aldrich); and 1 g/L L-glutamic acid (Sigma-Aldrich). The pH of the MIX6 medium was adjusted to 7.9 using 10 M NaOH (Sigma-Aldrich), and the medium was sterilized by filtration through a 0.22 μm PTFE filter (Corning).

### Preparation of *S. aureus* cells expressing fluorescent proteins

The plasmid pCN51F was prepared by cleaving the plasmid pCN51^60^ with the restriction enzymes SphI and NarI (New England Biolabs) and ligating it with annealed oligonucleotides carrying a Cerulean terminator. This insertion led to removing used restriction sites and introducing new NarI and Bg1II sites. The coding sequence of the red fluorescent protein mCherry2-L^36^ was codon-optimized for *S. aureus*, commercially synthesized, and inserted into the EcoRV site of pUC57-Amp (GeneWiz). The obtained plasmid was cleaved with SphI/NarI and ligated with annealed oligonucleotides carrying the constitutive capA promoter. After the transformation, constructs were cleaved with NarI/Bb1II and ligated into pCN51F, yielding reporter plasmids. These plasmids were then electroporated into *S. aureus* ISP479C rsbU+.^61^

### Preparation of *P. aeruginosa* cells expressing fluorescent proteins

The plasmid pSEVA238-msfGFP2x-KAN182 ^44^ was digested using PshAI/SwaI (NEB) restriction enzymes, treated with Shrimp Alkaline Phosphatase (NEB) and purified by agarose gel electrophoresis. Ligation with gentamycin cassette PCR amplified from the plasmid pACEBac1 (GenevaBiotech) was performed O/N at 16 °C using T4 DNA Ligase (NEB). Competent *E. coli* NEB 5-alpha (NEB) cells stored at −80 °C were thawed on ice and transformed with the ligation mixture according to the manufacturer’s protocol. The colonies were screened for the presence of the desired plasmid using colony PCR. The correctness of the construct was verified by sequencing. The prepared plasmid pSEVA238-msfGFP-GENT (∼100 ng) was electroporated into electroporation-competent *P. aeruginosa* BAA-28 cells. The cells were plated after 4 hours of vigorous shaking at 37 °C on an LB agar plate with gentamycin (10 µg/ml).

### Cultivation of biofilms in LSFM imaging chamber

Overnight colonies of *S. aureus* were resuspended and diluted to OD_600_ 0.01 in MIX6 medium containing 50 μg/ml erythromycin (Sigma-Aldrich), 1 μM POPO-1 (Thermo Fisher Scientific), and 2.5 μg/ml WGA Alexa Fluor 488 (Thermo Fisher Scientific). Overnight colonies of *P. aeruginosa* were resuspended and diluted to OD_600_ 0.05 or 0.01 in MIX6 medium containing 10 μg/ml gentamycin (Sigma-Aldrich), 0.5 μM BOBO-3 (Thermo Fisher Scientific), and 1 mM 3-mBz (m-toluic acid, Sigma-Aldrich) to induce msfGFP production. The inoculum was injected into an FEP tube (Zeus) with an inner diameter of 300 μm and placed in an LSFM microscope chamber filled with water kept at 37°C. After 4 hours of incubation, the unattached cells were washed away by pumping MIX6 medium through the tube. Biofilm growth was imaged every 30 min.

### Preparation of conditioned media

*S. aureus* and *P. aeruginosa* cultures (10 ml each) with an initial OD_600_ of both species of 0.01 were seeded in MIX6 medium in a 100 mm culture dish (Sigma-Aldrich) for 24 hours at 37°C without shaking. After 24 hours, the culture supernatant was collected, centrifuged at 5000 x g, passed through a 0.22 μm PTFE filter (Corning), and directly used for LSFM experiments or LC-MS/MS analysis.

### LC-MS/MS analysis of HQNO and other metabolites

Ethyl acetate (1 ml) was added to 0,5 ml of medium from bacterial culture, and the mixture was vortexed for 1 minute. The organic phase was collected, and the ethyl acetate was evaporated using a nitrogen gas stream. The dried sample was reconstituted in 160 μl of methanol, and 5 μl of the extract was injected into a liquid chromatographic column at a flow rate of 0.4 ml/min (Agilent) using an Ascentis Express-90 A Phenyl-Hexyl column (4.6 x 150 mm, 2.7 µm) (Supelco) maintained at 45°C. The HPLC separation system used 0.1% formic acid in water as the mobile phase A and 0.1% formic acid in methanol as the mobile phase B. The gradient profile was as follows: isocratic for 9 min, a linear gradient from 80% to 100% B over 1 min, followed by 100% B for 6 min. The column was re-equilibrated for 22 min.

All mass spectrometry experiments were performed using a 6410 Triple Quad LC/MS system (Agilent, USA). Positive electrospray ionization mode (ESI+) and multi-reaction monitoring (MRM) were used to identify and quantify selected analytes. HQNO (2-heptyl-4-hydroxyquinoline N-oxide) was quantified using authentic standard (Cayman Chemical Company), and other metabolites were quantified using the response factor of HQNO. Collision energy was 34 V for all compounds. Comparisons of the amounts of metabolites released into the medium were made using a *post-hoc* Tukey test. A *P* value of <0.05 was considered significant.

### Light sheet fluorescence microscopy data collection and volume reconstruction

A Zeiss Lightsheet Z.1 microscope fitted with two illumination objectives (10x/0.2) and a Plan-apochromatic detection objective (20×/1.0 water immersion, WD 2,4 mm) and PCO edge 5.5 sCMOS camera was used for imaging biofilm development. The excitation laser wavelengths were 405 nm, 488 nm, and 561 nm. The laser intensities were kept within the 0.5-2% range. The light sheet thickness at its focal point was 3.80 μm. Images were taken every 30 min with a slice spacing of 0.53 μm. The size of the acquisition area was 372.7 μm x 372.7 μm. Data acquisition, processing, and visualization were performed using the analysis software Zen Blue 3.6 (Carl Zeiss AG). For multidirectional imaging, the samples were imaged at relative orientations of 0°, 30°, 60°, 90°, 120°, and 150°. A movie file was exported from multidimensional datasets (multichannel, Z-stack, time series, multiple orientations). Three-dimensional visualizations of the data were generated using the BigStitcher 1.2.10. plugin for Fiji and displayed using Fiji or ChimeraX 1.4.^34, 35, 62^

### Confocal microscopy

The static distribution of biofilm matrix components in *S. aureus* biofilm was imaged using a Zeiss LSM 880 laser scanning confocal microscope with AiryscanFast super-resolution equipped with a C-Apochromat 63x/1.20 W objective. The excitation laser wavelengths were 405 nm, 488 nm, and 561 nm. Z-stacks were recorded with 0.27 μm intervals. The image data were processed using the software Zen Black (Carl Zeiss AG).

### Radial profile plot

The Intensity of signals from 3D biofilm reconstructions was analyzed using the Radial profile plot plugin for Fiji from Paul Baggethun. This method produces a profile plot of normalized integrated intensities as a function of distance from a point in the image. The radial profile plots of intensities were made for the z-projection of the 3D reconstructions of the whole FEP tube and for selected parts containing biofilm clusters. The profile was plotted, including the standard deviation of the values.

### Growth curves of S. aureus and P. aeruginosa

Overnight colonies of *S. aureus* mCherry2-L ISP479C rsbU+ or *P. aeruginosa* msfGFP BAA-28 were diluted in MIX6 medium to OD_600_ 0.01. Suspensions were pipetted in triplicates onto a 96-well cell culture microplate (SPL Life Sciences) with a final volume of 200 µl, and incubated at 37°C with constant shaking. Bacterial growth was monitored by measuring OD_600_ every 30 minutes for 20 hours using a Tecan Infinite M200 microplate reader (Tecan).

### Determination of emulsification properties of a conditioned medium

Two milliliters of sunflower oil were added to 2 ml of conditioned medium and vortexed at 3200 RPM for 2 min. The mixture was allowed to stand for 10 min, and the presence of emulsion was detected.

## Supplementary movie legends

**Supplementary movie 1. Time-lapse of *S. aureus* biofilm development.** *S. aureus* cells are shown in orange, eDNA in blue, and PIA in green. Images were taken every 30 minutes for 48 hours. The movie was prepared from a time-lapse dataset using the analysis software Zen.

**Supplementary movie 2. Time-lapse of *P. aeruginosa* biofilm development.** *P. aeruginosa* cells are shown in green and eDNA in blue. Images were taken every 30 minutes for 48 hours. The movie was prepared from a time-lapse dataset using the analysis software Zen.

**Supplementary movie 3. Time-lapse of *P. aeruginosa* and *S. aureus* biofilm development with equal OD_600_ ratio.** *P. aeruginosa* cells are shown in green and *S. aureus* cells in orange. The inoculation OD_600_ for both strains was 0.01. Images were taken every 30 minutes for 40 hours. The movie was prepared from a time-lapse dataset using the analysis software Zen.

**Supplementary movie 4. Time-lapse of *P. aeruginosa* and *S. aureus* biofilm development with higher inoculation OD_600_ in favor of P. aeruginosa.** *P. aeruginosa* cells are shown in green and *S. aureus* cells in orange. The inoculation OD_600_ for *P. aeruginosa* was 0.05, and that of *S. aureus* was 0.01. Images were taken every 30 minutes for 40 hours. The movie was prepared from a time-lapse dataset using the analysis software Zen.

**Supplementary movie 5. Effect of addition of conditioned medium to *S. aureus* biofilm.** *S. aureus* cells are shown in orange, eDNA in blue, and PIA in green. Conditioned medium (blue background fluorescence) obtained from coculture cultivation was added to a *S. aureus* biofilm 19-20 hours post-inoculation. We used concentrated conditioned medium or conditioned medium diluted 1:1 with MIX6. Images were taken every 30 minutes until biofilm dispersion. The movie was prepared from a time-lapse dataset using the analysis software Zen.

**Supplementary movie 6. Effect of HQNO on *S. aureus* biofilm.** *S. aureus* cells are shown in orange, eDNA in blue, and PIA in green. HQNO (150 ng/ml) was added to the *S. aureus* biofilm after 20 h of growth. Images were taken every 30 minutes until biofilm dispersion. The movie was prepared from a time-lapse dataset using the analysis software Zen.

## Supplementary figures

**Supplementary Fig. 1.**
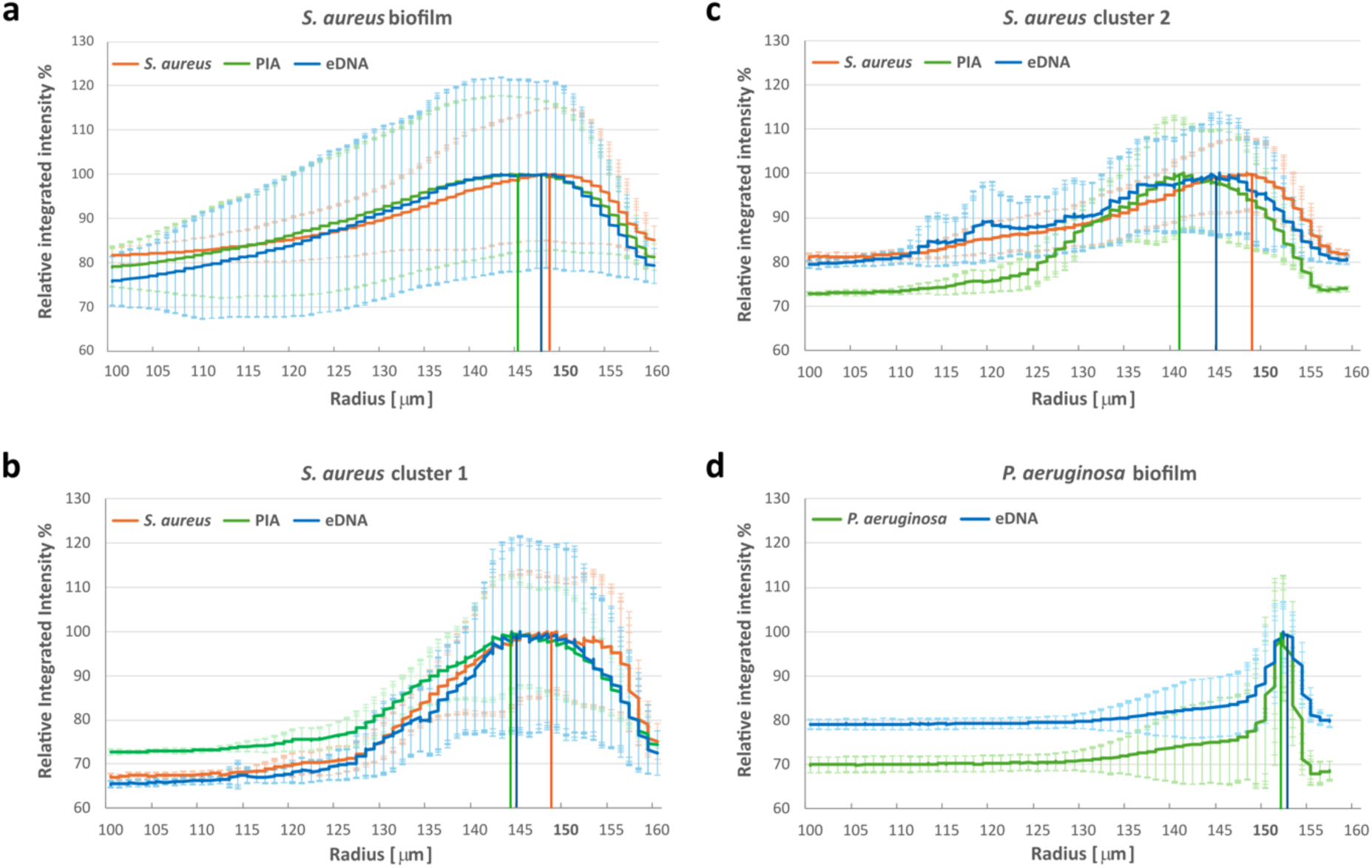
Radial profile plots of biofilm fluorescence intensities. Radially integrated intensities of *S. aureus* biofilm fluorescence calculated on a relative scale for the whole FEP tube **(a),** cluster 1 **(b),** and cluster 2 **(c)** from Fig. 7. Radially integrated intensities of *P. aeruginosa* biofilm fluorescence on a relative scale calculated for the whole FEP tube **(d).** Maximum intensities of the individual fluorescence channels are marked with their respective colors. The x-axis corresponds to the radius of the FEP tube, where 0 (not shown) is at the center of the tube and 150 μm corresponds to the inner wall of the tube.

**Supplementary Fig. 2.**
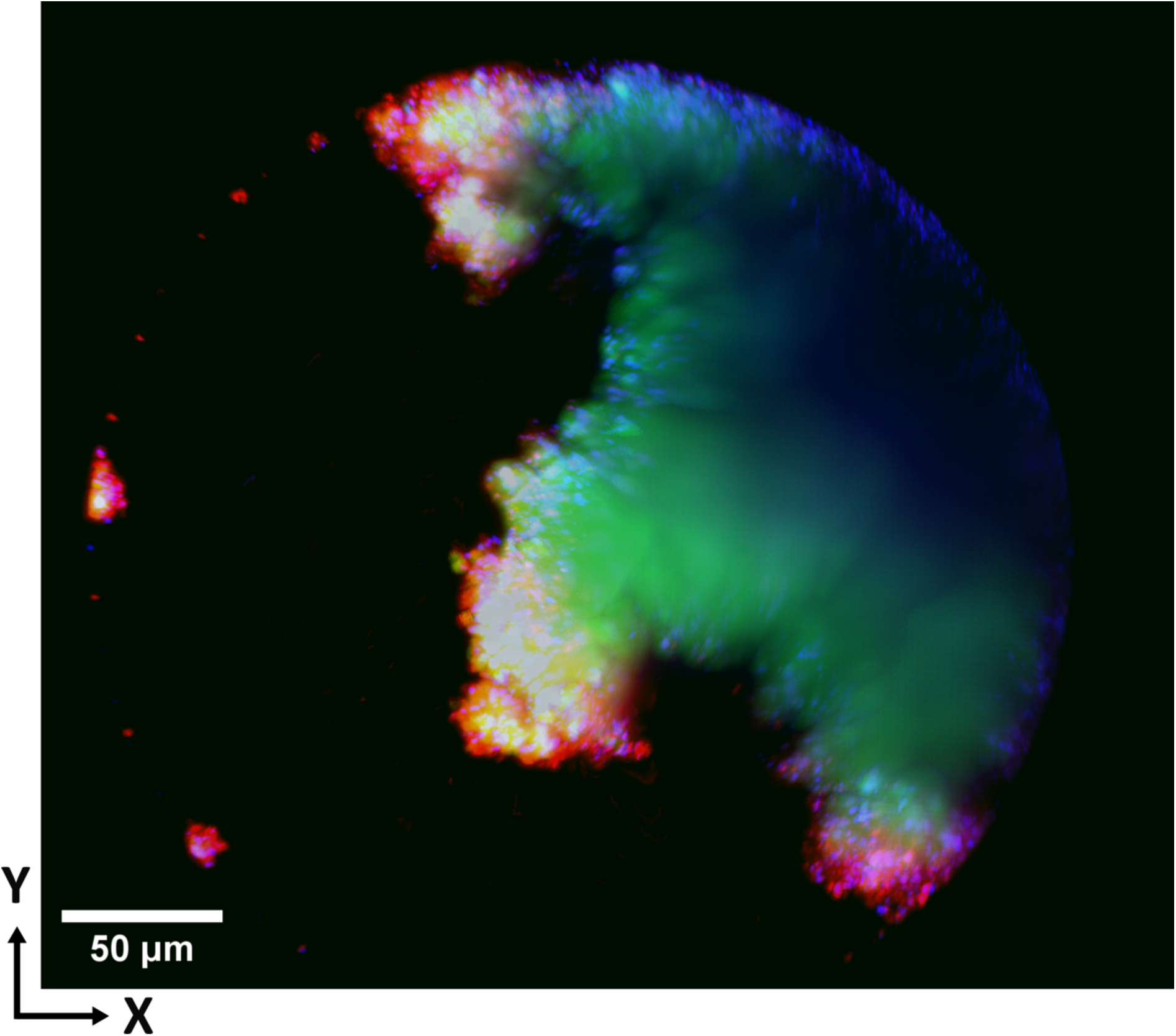
Three-dimensional reconstruction of five days old *S. aureus* biofilm. Metabolically active *S. aureus* cells are at the biofilm surface. *S. aureus* cells are shown in red, eDNA in blue, and PIA in green.

**Supplementary Fig. 3.**
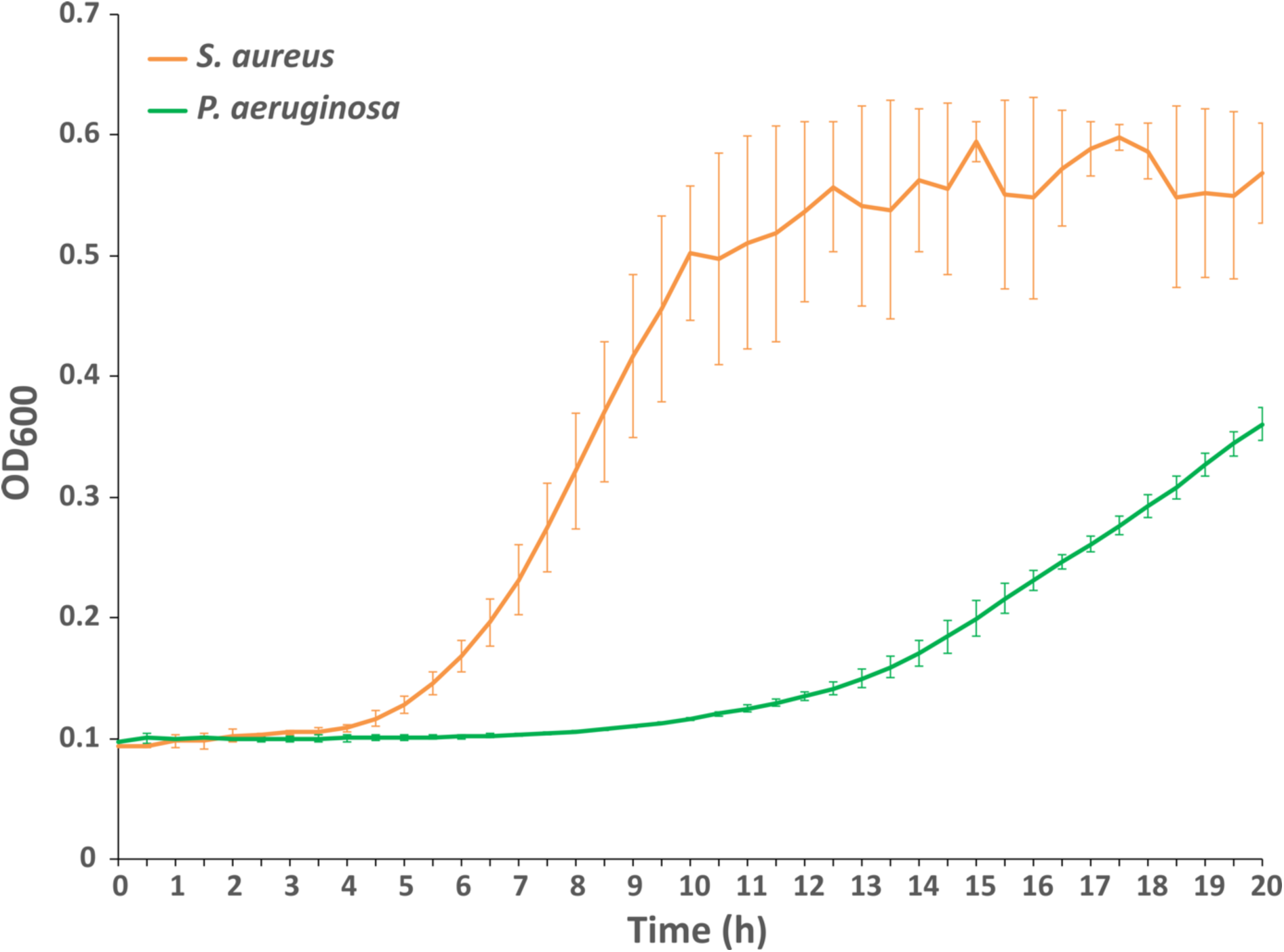
Plot of *S. aureus* and *P. aeruginosa* growth kinetics in MIX6 medium. *S. aureus* mCherry2-L ISP479C rsbU+ or *P. aeruginosa* msfGFP BAA-28 were diluted in MIX6 medium to OD_600_ 0.01. OD_600_ was monitored every 30 minutes for 20 hours.

**Supplementary Fig. 4.**
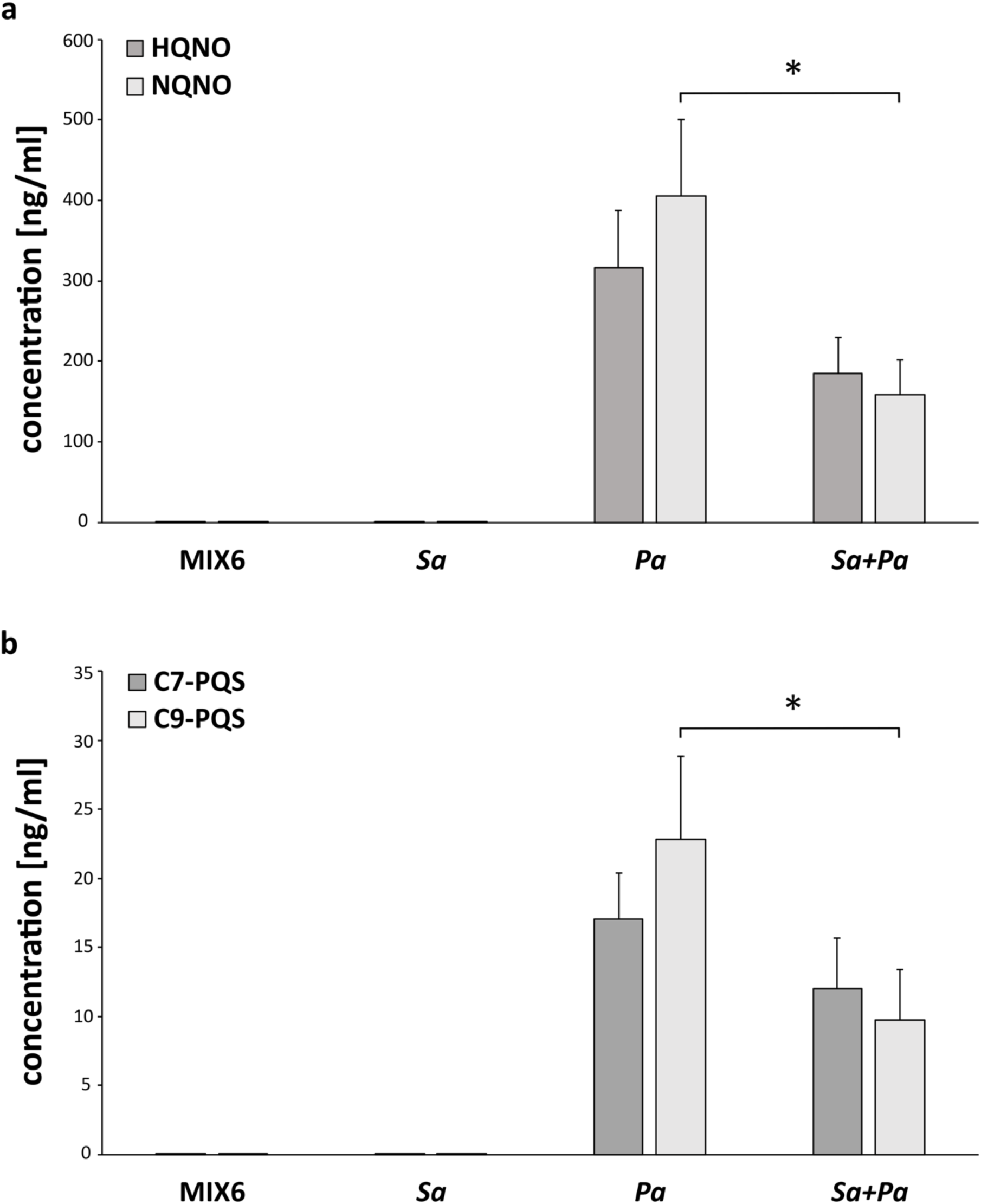
LC-MS/MS analysis of respiratory toxins secreted by *P. aeruginosa*. **(a)** Identification of 2-heptyl-4-hydroxyquinoline N-oxide (HQNO) and 2-nonyl-4-hydroxyquinoline N-oxide (NQNO). **(b)** Identification of 2-heptyl-3-hydroxy-4(1H)-quinolone (C7-*Pseudomonas* quinolone signal) and 2-hydroxy-2-nonyl-4(1H)-quinolone (C9-*Pseudomonas* quinolone signal) released into the medium after 24 hours of static cultivation of *S. aureus (Sa)*, *P. aeruginosa (Pa)* or a coculture biofilm *(Sa+Pa)*. MIX6 medium was used as a negative control. The results are expressed as means ± standard deviation of three independent experiments. The symbol * indicates a statistically significant difference between *P. aeruginosa* and coculture (P<0.05).

**Supplementary Fig. 5.**
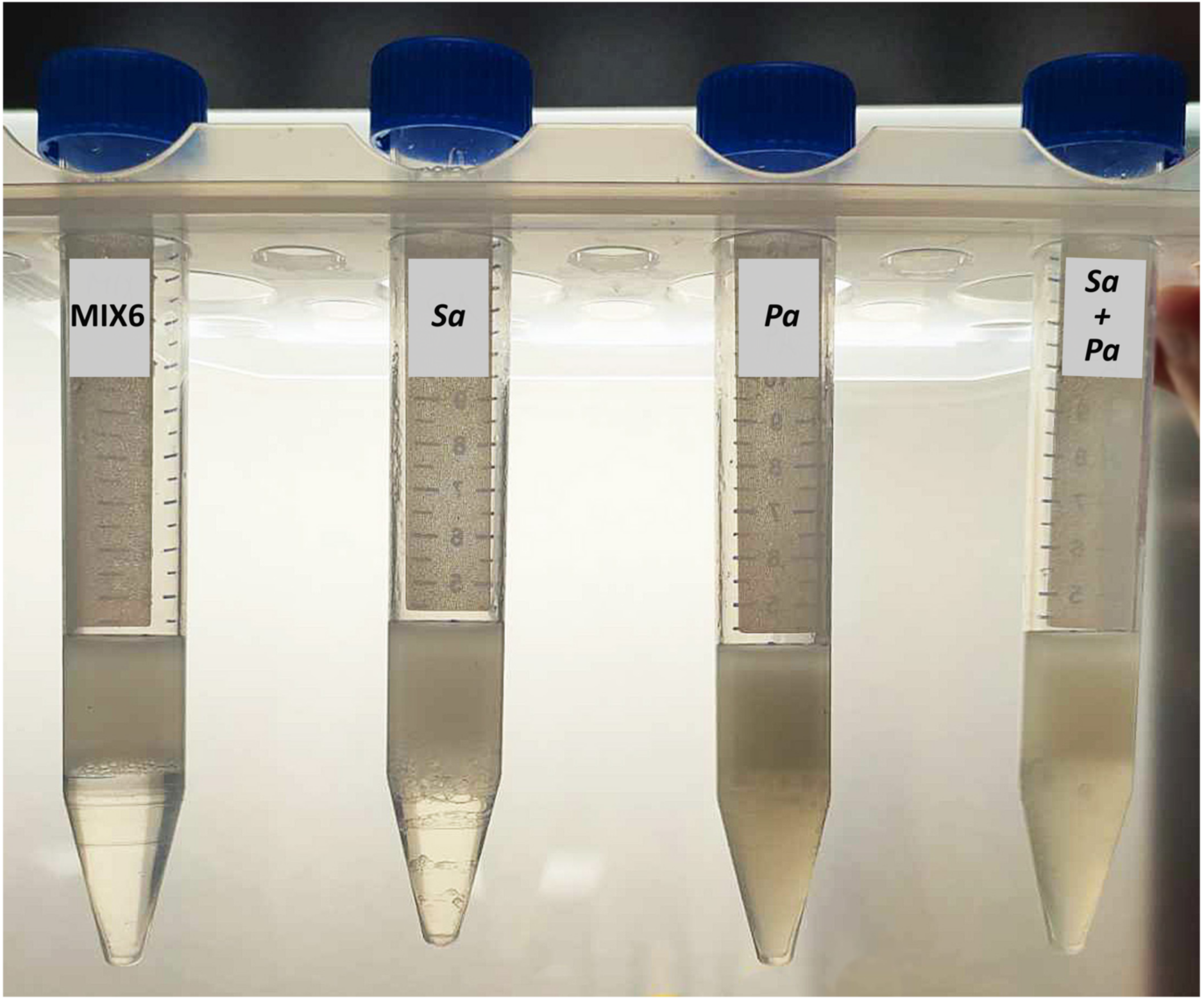
Determination of rhamnolipids production by biofilms using emulsification test. The emulsification activity of solutions collected from bacterial cultures was determined using sunflower oil. MIX6 medium was used as a negative control. Media from *S. aureus (Sa), P. aeruginosa (Pa)*, and a coculture biofilm *(Sa+Pa)* after 24 hours of static cultivation.

**Table S1:** List of bacterial strains and plasmids used.

| bacterial strain | resistance | purpose | origin | reference |
| --- | --- | --- | --- | --- |
| *E. coli* NEB 5-α | | Molecular cloning | New England Biolabs | |
| *S. aureus* ISP479C<br>rsbU+ |  | Biofilm<br>experiments | Prof. Lasa, University<br>of Navarra, Spain | Sullam et<br>al., 1996 |
| *P. aeruginosa* BAA-28 |  | Biofilm<br>experiments | ATCC, University<br>Boulevard Manassas;<br>USA |  |
| plasmids |  |  |  |  |
| pUC57-Amp-<br>mCherry2-L | Amp | Gene-synthetized<br>mCherry2-L | GeneWiz |  |
| pCN51 | Amp/Ery | Cloning vector | GeneWiz | Charpentier<br>et al., 2004 |
| pCN51-CapA-mCherry2-L | Amp/Ery | mCherry2-L<br>reporter plasmid | this study | Balleza et.<br>al., 2018 |
| pACEBac1 | Gent | Source of<br>gentamycin<br>cassette | GenevaBiotech |  |
| pSEVA238_msfGFP2x<br>_Km | Kan | msfGFP reporter<br>plasmid | SEVA | Calles et al.,<br>2019 |
| pSEVA238_msfGFP2x<br>_Gm | Gent | msfGFP<br>reporter plasmid | this study |  |

